# Antifungal efficacy of natural antiseptic products against *Candida auris*

**DOI:** 10.1101/2023.09.22.559057

**Authors:** Wing-Gi Wu, Kristine Shik Luk, Mei-Fan Hung, Wing-Yi Tsang, Kin-Ping Lee, Bosco Hoi-Shiu Lam, Ka-Lam Cheng, Wing-Sze Cheung, Hau-Ling Tang, Wing-Kin To

**Author notes:** Correspondence to: Dr Wing-Gi WU, Department of Pathology, Princess Margaret Hospital, 2-10 Princess Margaret Hospital Road, Lai Chi Kok, Kowloon, Hong Kong.

## Abstract

*Candida auris* is an emerging fungal pathogen responsible for healthcare associated infections and outbreaks with high mortality around the world. It readily colonizes the skin, nares, respiratory and urinary tract of hospitalized patients, and such colonization may lead to invasive *Candida* infection in susceptible patients. However, there is no recommended decolonization protocol for *C. auris* by international health authorities. The aim of this study is to evaluate the susceptibility of *C. auris* to commonly used synthetic and natural antiseptic products using an in vitro, broth microdilution assay. Synthetic antiseptics including chlorhexidine, povidone-iodine, and nystatin were shown to be fungicidal against *C. auris*. Among the natural antiseptics tested, tea tree oil and manuka oil were both fungicidal against *C. auris* at concentrations less than or equal to 1.25% (v/v). Manuka honey inhibited *C. auris* at 25% (v/v) concentrations. Among the commercial products tested, manuka body wash and mouthwash were fungicidal against *C. auris* at concentrations less than or equal to 0.39% (w/v) and 6.25% (v/v) of products as supplied for use, respectively, while tea tree body wash and Medihoney^TM^ wound gel demonstrated fungistatic properties. In conclusion, this study demonstrated good in vitro antifungal efficacy of tea tree oil, manuka oil, manuka honey, and commercially available antiseptic products containing these active ingredients. Future studies are warranted to evaluate the effectiveness of these antiseptic products in clinical settings.

## INTRODUCTION

The fungal pathogen *Candida auris* was first described in 2009 after isolation from the ear of a patient in Japan [1]. It has since been reported in more than 40 countries across six continents, causing a wide range of healthcare-associated outbreaks and infections [2], with the mortality rate varying from 30 to 60%, due to the high virulence and resistance to several antifungals [3]. The recent inaugural World Health Organization fungal priority pathogen list highlights the urgent threat of *C. auris* to global public health [4]. With its biofilm-forming ability and tolerance to high temperature and salt concentrations, *C. auris* can persist and survive in the human skin and environmental surfaces for several months following contact with colonized patients [5–7]. This provides additional opportunities for transmission and colonization within the healthcare settings. Furthermore, the minimum time to acquire *C. auris* from a patient or their immediate environment is 4 hours or less, reinforcing the importance of vigilant infection control measures [8]. This fungal superbug can also resist certain disinfectants and is well adapted to healthcare environments [9].

In 2019, a cluster of *C. auris* colonization affecting 15 patients was reported in Princess Margaret Hospital (PMH), Kowloon West (KW) cluster, Hong Kong. Whole genome sequencing identified the strains as Clade I/South Asia, associated with ERG11 mutation conferring azole resistance [10]. The clade is also known to cause invasive infections and outbreaks frequently [6]. Four regional hospitals (a total of 3,878 beds) serve an estimated population of 1.4 million in the KW cluster. In spite of enhanced surveillance and infection control measures, *C. auris* cases and transmission have continued to increase due to pandemic-related strain on the healthcare system. As of April 2023, there were 364 cases of *C. auris* reported in KW Cluster, Hong Kong.

*C. auris* readily colonizes the axillae, groin, nares, respiratory, and urinary tract of hospitalized patients [6]. Compared to other *Candida* species, gastrointestinal tract colonization is less frequent for *C. auris*. [11]. Colonization predisposes to invasive infection, with mechanical ventilation and placement of invasive devices identified as major risk factors [6]. A descriptive analysis of *C. auris* outbreaks in New York State reported that 4% of *C. auris* colonized patients developed bloodstream infection, with a rate of 0.3 per 1,000 patient-days [12]. These studies provided evidence that an important strategy for preventing severe candidiasis is the elimination of *C. auris* carriage to preclude subsequent risk of endogenous infection. Nevertheless, there are no well-established guidelines for *C. auris* decolonization. For this reason, more evidence from in vitro antifungal susceptibility testing of topical antiseptics against *C. auris* is needed to provide the basis of decolonization regimen before clinical trials. Synthetic antiseptics including chlorhexidine, povidone-iodine, isopropyl alcohol, and nystatin have been most studied for their antifungal activities against *C. auris* in vitro [13–15]. While being one of the commonly used disinfectants in healthcare settings, chlorhexidine has been shown to have variable effectiveness for *C. auris* depending on the formulation and in vitro testing methods. *Sherry et al.* and *Abdolrasouli et al.* showed chlorhexidine was effective for *C. auris* using broth dilution antifungal susceptibility testing [16–17]. *Moore et al*. reported that 2% chlorhexidine solution with isopropyl alcohol formation was effective in killing *C. auris* with a 2-min contact time, while chlorhexidine alone without isopropyl alcohol failed to kill *C. auris* [13].

By far, most clinical experience of *C. auris* decolonization was reported in outbreak settings as part of the infection control measures. *Schelenz et al.* first reported a decolonization regimen using a five-day protocol with twice-daily 2% chlorhexidine body washes, 0.2% chlorhexidine mouthwash, and oral nystatin during hospital outbreak management [8]. Based on this early experience, decolonization on our *C. auris* colonized cohorts using a protocol of twice-daily chlorhexidine bath, chlorhexidine mouthwash and povidone iodine applied to axilla, groin and nares was attempted as an outbreak control measure. Frequent chlorhexidine baths were poorly tolerated. More importantly, none of our patients were successfully decolonized (0/14 patients; unpublished). In addition, subsequent studies have reported poor clinical efficacy of chlorhexidine-based regimens for decolonizing *C. auris,* with high rates of persistent colonization [9]. A murine model study demonstrated that *C. auris* could enter the dermis without causing overt histopathologic signs of inflammation, resulting in deeper skin layer residence [18]. Another study based on ex vivo human skin model has illustrated chlorhexidine had limited permeation into deeper skin layers [19], and this might contribute to persistent *C. auris* colonization despite its in vitro efficacy.

The limited success of traditional decolonization using chlorhexidine has highlighted the need to explore alternative regimens and approaches. Natural substances with antimicrobial properties are potentially good candidates as their antimicrobial effects are based on multiple compounds and would not mount drug-selecting pressure. They are also well tolerated as skin washes in general. *Fernades et al.* have shown that tea tree (*Melalenca alternifolia*) oil had good in vitro antifungal activity against *C. auris* in both planktonic cells and biofilms [20]. *Johnson et al.* have also shown that the active components of tea tree oil could augment the activity of chlorhexidine for *C. auris* decolonization in the porcine skin model [21]. Another potential natural antiseptic is manuka (*Leptospermum scoparium*), which is also known as the “New Zealand tea tree”. It has been a part of the traditional Māori medicine with fungicidal properties towards *C. albicans* and *C. glabrata* [22]. Notably, manuka honey was found to be effective in inhibiting various *Candida* species [23]. There are multiple antimicrobial constituents of the natural extracts, namely monoterpenes and sesquiterpenes (tea tree and manuka oil), triketones (manuka oil), methylglyoxal, polyphenolic compounds and bee defensin-1 (manuka honey) [24–26]. Therefore, unlike other synthetic antimicrobials, there have been no reports of clinical isolates with acquired resistance to the natural antiseptics [27–28]. Clinical trials have demonstrated the efficacy of manuka honey and tea tree oil in the treatment of oral candidiasis, vulvovaginal candidiasis and successful decolonization for methicillin-resistant *Staphylococcus aureus* (MRSA) carriers [29–34], making them attractive alternatives against *C. auris*. These natural components are also widely used as active ingredients for their antiseptic effects in commercial hygiene products such as body wash, mouthwash and wound gel. To pave the way for evaluating the clinical application of the natural antiseptics against *C. auris*, this study aimed to determine the in vitro antifungal susceptibility of *C. auris* to the natural antiseptics and their commercial products.

## MATERIALS AND METHODS

### Characterization of *Candida auris* isolates

A total of 40 *C. auris* isolates belonging to clade I/South Asia (unpublished data) were obtained in the KW Cluster of Hong Kong from 2019 to 2022. All tested isolates were identified by matrix-assisted laser desorption ionization-time of flight mass spectrometry (Bruker Daltonics; Database version: IVD-9468 MSP) [35]. Reference strains *C. auris* JCM 15448 (clade II/East Asia), *C. auris* ATCC MYA-5002 (clade III/South Africa), *C. auris* ATCC MYA-5003 (clade IV/South America), *C. albicans* ATCC 90028, *C. glabrata* ATCC MYA-2950, *C. krusei* ATCC 6258, and *C. parapsilosis* ATCC 22019 were included. The study protocol was approved by the Research Ethics Committee of the Kowloon West Cluster, Hospital Authority, Hong Kong (KW/EX-21-098(160-14)).

### Antiseptics and preparations

Our experiment studied three groups of topical antiseptics. The first group included synthetic antiseptics products commonly used in healthcare settings: chlorhexidine gluconate (Microshield^®^ 4%, Australia), povidone-iodine (Betadine^®^ 10%, Turkey) and nystatin (100,000 units/mL, Pharmascience, Canada). The second group comprised natural antiseptics: tea tree oil (15%, The Body Shop, UK; 100% Kiwi Manuka^®^ , New Zealand), manuka oil (100%, Manuka Lab^®^ , New Zealand; 100%, Melora^®^ , New Zealand), manuka honey (UMF 15+ and 20+, Comvita^®^ , New Zealand; UMF 15+ and 20+, Melora^®^ , New Zealand), manuka hydrosol (Bioactives^®^ , New Zealand). The third group consisted of commercial antiseptic products containing active components from tea tree oil and/or manuka derivatives: manuka body wash (Manuka Lab^®^ , New Zealand), tea tree clearing body wash (The Body Shop^®^ , UK), manuka mouthwash (Manuka Lab^®^ , New Zealand), and Medihoney^TM^ antibacterial wound gel (Comvita^®^ , New Zealand). Dimethyl sulfoxide (DMSO, Sigma-Aldrich, UK) at 2% was added to water-insoluble products, including tea tree oil, manuka oil, antibacterial wound gel, and nystatin to enhance solubility in RPMI 1640 medium (Sigma, Paisley, UK). The effects of 2% DMSO in growth control was also studied to confirm no antifungal activity was added by the solvent.

### In vitro antifungal susceptibility of antiseptics

In vitro antifungal susceptibility of antiseptics was performed by broth microdilution method according to CLSI M27-A4 [36]. Briefly, 100 µL of RPMI 1640 medium with different concentrations of the antiseptics were added to 96-well microtiter plates (Saining Biotechnology, China). *Candida* isolates were suspended in 2 mL sterile physiological saline to obtain 0.5 McFarland standard suspension using the DensiCHEK Plus instrument (BioMerieux, France), corresponding to 1-5 x 10^6^ cells per mL, then further diluted with RPMI 1640 medium to obtain a 1:1000 final inoculum containing 1-5 x 10^3^ cells per mL. The yeast suspensions (100 µL) were added to the 96-well plate with the product suspensions (100 µL) and incubated aerobically at 35°C for 24 h. The minimum inhibitory concentration (MIC) endpoint was determined visually and defined as the lowest concentration of a product that caused a 50% decrease in growth (prominent decrease in turbidity) relative to the growth in the control well. To assess the minimal fungicidal concentration (MFC) endpoint, the broth was first assessed for any visible growth. For those without visible growth, 100 µL of broth was taken from the wells and inoculated onto Sabouraud dextrose agar (BD DIFCOTM, USA). Following incubation at 35°C for 48 h, the colony-forming unit (CFU) was counted to assess viability. The antiseptic was defined as fungicidal if there was no growth on inoculated plates, whereas it was defined as fungistatic if it inhibited growth but not fungicidal. Antifungal was defined as a product with either fungistatic or fungicidal effects. Each isolate was tested in duplicate separately and the assay was repeated if the MIC or MFC differed by more than one dilution. Antifungal drug susceptibility of *C. krusei* ATCC 6258 and *C. parapsilosis* ATCC 22019 was performed in each run of broth microdilution as controls [37].

For antiseptics demonstrating fungistatic effects on clinical *C. auris* isolates, log growth reduction in CFU compared to the growth control (yeast suspension in RPMI) after 24-hour incubation was measured, and the test was repeated in triplicate. The yeast suspension preparation and incubation procedures were the same as the aforementioned section. Mann-Whitney U test was used to analyze the significance of difference in CFU count compared to the growth control, and one-way ANOVA, Kruskal-Wallis, or Mann-Whitney U tests were used to compare the log growth reduction in CFU by the antiseptics at different concentrations. Pairwise comparisons were performed using Bonferroni or Dunn’s test when suitable. P value <0.05 was considered statistically significant. All statistical analyses were performed using STATA18.0 (StataCorp. 2023. Stata Statistical Software: Release 18. College Station, TX: StataCorp LLC.).

### In vitro antifungal susceptibility testing of antifungal drugs

To confirm the local resistance pattern of our isolates to conventional antifungal drugs, in vitro susceptibility was determined using Sensititre^TM^ YeastONE^TM^ YO10 AST Plate (ThermoFisher Scientific, UK). The antifungal drugs included anidulafungin, micafungin, caspofungin, 5-flucytosine, posaconazole, voriconazole, itraconazole, fluconazole and amphotericin B. The test was performed according to the procedures described by the manufacturer [38]. Centres for Disease Control and Prevention (CDC)’s tentative breakpoints for *C. auris* were used as a reference [39].

## RESULTS

Table 1 summarizes the antifungal activities of topical antiseptics against *Candida auris*. For the synthetic antiseptics, chlorhexidine, povidone-iodine and nystatin were fungicidal against *C. auris* at concentrations less than or equal to 21.99 mg/L (0.004%), 312.50 mg/L (0.313%), and 9.77 μg/mL (0.049%), respectively. The MICs and MFCs of chlorhexidine against *C. auris* strains, including the JCM 15448, were relatively higher when compared with that against *C. krusei* ATCC 6258, *C. parapsilosis* ATCC 22019, *C. albicans* ATCC 90028, and *C. glabrata* ATCC MYA-2950.

**Table 1:** In vitro susceptibility of *Candida* isolates to synthetic antiseptics, natural antiseptics and their commercial products

|  | Clinical *Candida auris* isolates (N=40)<br>(clade I/South Asia) |  |  |  | *Candida auris*<br>JCM 15448<br>(clade II/East Asia) |  | *Candida auris*<br>ATCC MYA 5002<br>(clade III/South Africa) |  | *Candida auris*<br>ATCC MYA 5003<br>(clade IV/South America) |  | *Candida krusei*<br>ATCC 6258 |  | *Candida parapsilosis*<br>ATCC 22019 |  | *Candida albicans*<br>ATCC 90028 |  | *Candida glabrata* ATCC<br>MYA-2950 |  |
| --- | --- | --- | --- | --- | --- | --- | --- | --- | --- | --- | --- | --- | --- | --- | --- | --- | --- | --- |
|  | MIC50 | MIC90 | MIC range | MFC range | MIC | MFC | MIC | MFC | MIC | MFC | MIC | MFC | MIC | MFC | MIC | MFC | MIC | MFC |
| **Synthetic antiseptics** |  |  |  |  |  |  |  |  |  |  |  |  |  |  |  |  |  |  |
| Chlorhexidine (%) | 0.001 | 0.001 | 0.0005-0.001 | 0.002-0.004 | 0.001 | 0.002 | 0.001 | 0.002 | 0.001 | 0.002 | 0.0002 | 0.0005 | 0.00006 | 0.0002 | 0.0005 | 0.001 | 0.0005 | 0.0005 |
| Povidone-iodine (%) | 0.078 | 0.078 | 0.039-0.078 | 0.078-0.313 | 0.039 | 0.16 | 0.078 | 0.156 | 0.078 | 0.156 | 0.039 | 0.31 | 0.078 | 0.625 | 0.16 | 0.31 | 0.16 | 0.31 |
| Nystatin (µg/mL) | 2.44 | 2.44 | 1.22-2.44 | 2.44-9.77 | 1.22 | 1.22 | 1.22 | 2.44 | 1.22 | 2.44 | 1.22 | 2.44 | 0.61 | 2.44 | 1.22 | 4.88 | 1.22 | 2.44 |
| **Natural antiseptics** |  |  |  |  |  |  |  |  |  |  |  |  |  |  |  |  |  |  |
| Manuka honey 15+ (Comvita) (v/v) (%) | 25 | 25 | 6.25-25 | >50 | 0.20 | >50 | 25 | >50 | 25 | >50 | 25 | >50 | 25 | >50 | 25 | >50 | 1.56 | >50 |
| Manuka honey 20+ (Comvita) (v/v) (%) | 25 | 25 | 3.13-25 | >50 | 0.20 | >50 | 25 | >50 | 25 | >50 | 12.5 | >50 | 25 | >50 | 25 | >50 | 1.56 | >50 |
| Manuka honey 15+ (Kiwi Manuka) (v/v) (%) | 25 | 25 | 3.13-25 | >50 | 0.20 | >50 | 25 | >50 | 25 | >50 | 12.5 | >50 | 25 | >50 | 25 | >50 | 1.56 | >50 |
| Manuka honey 20+ (Kiwi Manuka) (v/v) (%) | 25 | 25 | 6.25-25 | >50 | 0.20 | >50 | 25 | >50 | 25 | >50 | 12.5 | >50 | 25 | >50 | 25 | >50 | 1.56 | >50 |
| Tea tree oil 15% (Body Shop) (v/v) (%) | 0.004 | 0.004 | 0.004 | 2.08-4.17 | 0.004 | 1.04 | 0.001 | 0.31 | 0.001 | 0.31 | 0.002 | 2.08 | 0.004 | 4.17 | 0.039 | 0.63 | 0.16 | 0.63 |
| Tea tree oil 100% (Kiwi Manuka) (v/v) (%) | 0.31 | 0.31 | 0.31 | 0.63-1.25 | 0.31 | 0.63 | 0.31 | 1.25 | 0.31 | 1.25 | 0.63 | 1.25 | 0.63 | 1.25 | 0.63 | 1.25 | 0.63 | 1.25 |
| Manuka oil 100% (Manuka Lab) (v/v) (%) | 0.31 | 0.31 | 0.16-0.31 | 0.63 | 0.08 | 0.63 | 0.31 | 0.63 | 0.31 | 0.63 | 0.31 | 1.25 | 0.31 | 1.25 | 0.63 | 2.5 | 0.08 | 0.63 |
| Manuka oil 100% (Melora) (v/v) (%) | 0.31 | 0.31 | 0.31 | 0.63 | 0.04 | 0.31 | 0.31 | 1.25 | 0.31 | 1.25 | 0.31 | 1.25 | 0.31 | 1.25 | 1.25 | 5.0 | 0.08 | 0.63 |
| Manuka hydrosol (v/v) (%) | >50 | >50 | >50 | >50 | >50 | >50 | >50 | >50 | >50 | >50 | >50 | >50 | >50 | >50 | >50 | >50 | >50 | >50 |
| **Commercial products** |  |  |  |  |  |  |  |  |  |  |  |  |  |  |  |  |  |  |
| Manuka body wash (w/v) (%) | 0.012 | 0.012 | 0.006-0.012 | 0.05-0.10 | 0.006 | 0.024 | 0.024 | 0.39 | 0.012 | 0.098 | 0.006 | 0.05 | 0.006 | 3.13 | 0.024 | 0.098 | 0.049 | 1.56 |
| Tea tree body wash (w/v) (%) | 0.012 | 0.012 | 0.006-0.39 | ≥50 | 0.003 | 25 | 0.006 | 50 | 0.006 | 50 | 0.002 | 25 | 0.003 | 50 | 0.049 | 25 | 0.20 | >50 |
| MEDIHONEY wound gel (w/v) (%) | 3.13 | 3.13 | 1.56-6.25 | ≥25 | 0.20 | 12.5 | 3.13 | >37.5 | 6.25 | >37.5 | 0.39 | 25 | 0.20 | >37.5 | 6.25 | >37.5 | 0.098 | 37.5 |
| Manuka mouthwash (v/v) (%) | 0.39 | 0.39 | 0.20-0.39 | 3.13-6.25 | 0.39 | 6.25 | 0.78 | 3.13 | 1.56 | 3.13 | 0.10 | 6.25 | 0.20 | 6.25 | 1.56 | 6.25 | 1.56 | 6.25 |

For the natural antiseptics, whilst manuka honey exhibited fungistatic properties against clinical *C. auris* isolates at 25% (v/v), it was not fungicidal. Pure (equates to 100% in neat formula) tea tree and manuka oil were fungicidal against clinical *C. auris* isolates at concentrations less than or equal to 1.25% and 0.63% (v/v), respectively. Commercial products containing active ingredients of manuka honey, manuka oil and/or tea tree oil were shown to be active in vitro against *C. auris* and other *Candida* species. Manuka body wash and mouthwash were fungicidal against *C. auris* at concentrations less than or equal to 0.39% (w/v) and 6.25% (v/v), respectively. Manuka body wash, however, exhibited weaker antifungal activity against *C. parapsilosis* [MFC 3.13% (w/v)]. Interestingly, the MICs of manuka honey against *C. auris* JCM 15448 (clade II) were 128-fold lower than those against the clinical counterparts (clade I) and the two ATCC strains of clades III and IV. Tea tree body wash and Medihoney^TM^ wound gel even became fungicidal against *C. auris* JCM 15488 at concentrations of 25% (w/v) and 12.5% (w/v), respectively.

Manuka honey [25-50% (v/v)], tea tree body wash [25-50% (w/v)] and Medihoney^TM^ wound gel [25-37.5% (w/v)] demonstrated fungistatic properties against clinical *C. auris* isolates in vitro, but they were not fungicidal. They resulted in a statistically significant log reduction in CFU compared to the growth control (p <0.01). The growth control yielded around 1.5-5 x 10^7^ CFU/mL after 24 h incubation. At a concentration of 37.5%, tea tree body wash and the Medihoney^TM^ wound gel achieved >4 log reduction in CFU while manuka honey achieved 2 to 3 log reduction in CFU after 24 h of incubation, as shown in Figure 1. The inhibitory effects of manuka honey and tea tree body wash against *C. auris* were concentration-dependent (p <0.05). Medihoney^TM^ wound gel at 25% and 37.5% resulted in a similar reduction (6-log) in CFU.

**Figure 1:**
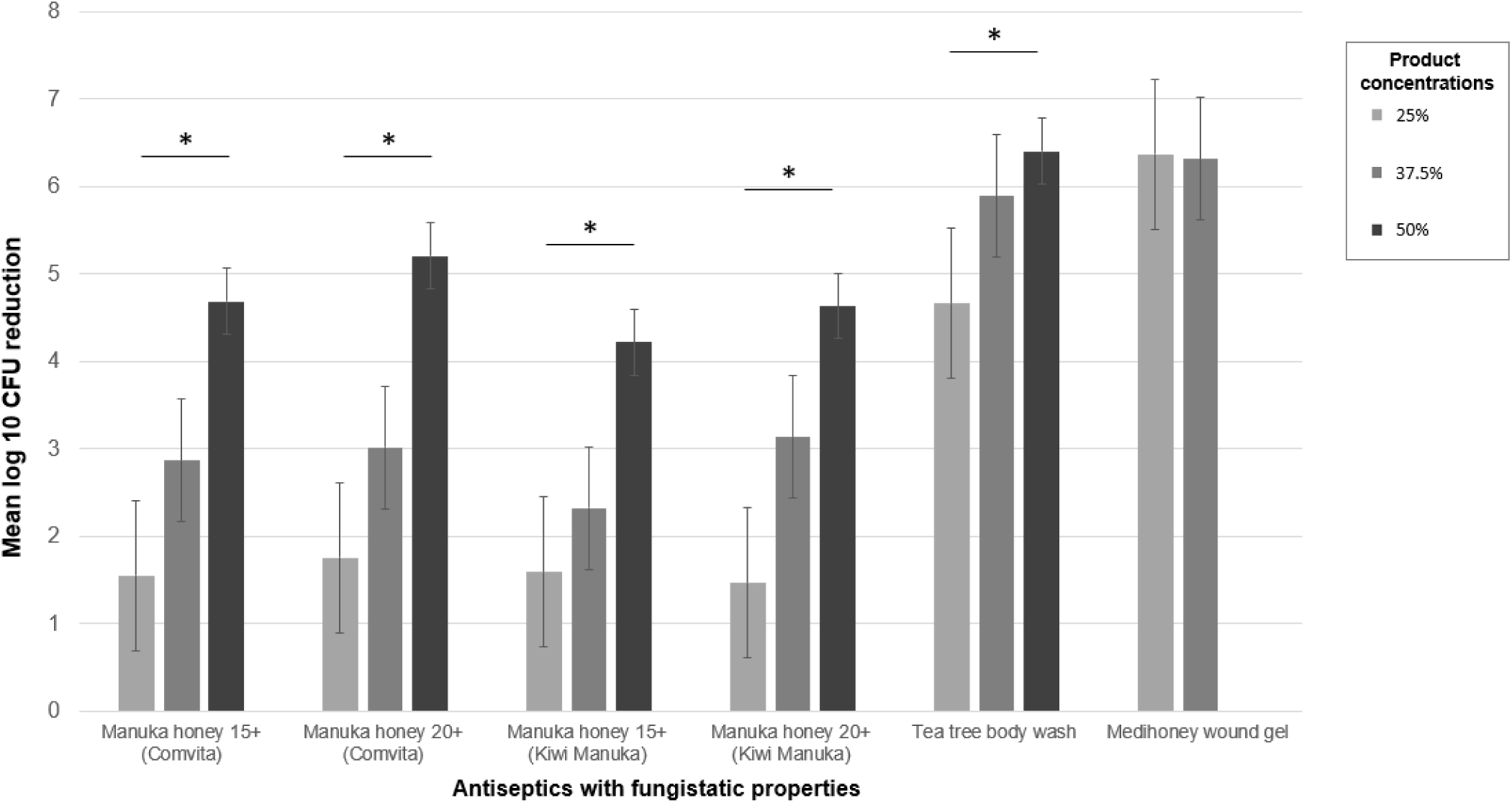
Concentration-dependent antifungal effect of natural antiseptics on *Candida auris*. *Candida auris* isolates (n = 3) were selected with MICs representative for MIC_90_ of the 40 isolates. The test products were diluted using RPMI to a test concentration of 25%, 37.5%, and 50% for manuka honey (v/v), tea tree body wash (w/v), and Medihoney^TM^ wound gel (w/v) (50% wound gel is not available for testing due to its insolubility at high concentrations). Log reductions in CFU compared with growth control (yeast suspension in RPMI) are shown. Significant differences (p < 0.05) between concentrations of antiseptics are indicated with an asterisk.

The antifungal susceptibility of common systemic antifungals against 40 clinical *C. auris* strains (clade I) was shown in table 2. All tested isolates (N=40) were resistant to fluconazole and 90% (N=36) were also resistant to Amphotericin B. Susceptibility to other azoles and echinocandins was variable.

**Table 2:** Antifungal drug susceptibility of clinical *Candida auris* isolates

|  | Clinical *C. auris* isolates (N=40) |  |  |  |
| --- | --- | --- | --- | --- |
|  | MIC<sub>50</sub> (mg/L) | MIC<sub>90</sub> (mg/L) | MIC range (mg/L) | Resistance, n (%) |
| **Amphotericin B** | 2 | 4 | 1-4 | 36 (90) |
| **Fluconazole** | 256 | >256 | 64 - >256 | 40 (100) |
| **Anidulafungin** | 0.25 | 0.25 | 0.12 - 8 | 1 (2.5) |
| **Micafungin** | 0.25 | 0.25 | 0.06 - 1 | 0 (0) |
| **Caspofungin** | 0.25 | 0.5 | 0.06 - >8 | 3 (7.5) |
| **Itraconazole** | 0.12 | 0.25 | 0.06 - >16 | NA |
| **Voriconazole** | 0.5 | 1 | 0.12 - >8 | NA |
| **Posaconazole** | 0.03 | 0.06 | 0.015 - >8 | NA |
| **5-flucytosine** | 0.12 | 0.12 | <0.06 - >64 | NA |

CDC's tentative nonsusceptible breakpoints: $\geq 2$ mg/L for amphotericin B, $\geq 32$ mg/L for fluconazole, $\geq 4$ mg/L for echinocandins anidulafungin and micafungin and $\geq 2$ mg/L for caspofungin. For the triazoles itraconazole, voriconazole and posaconazole, and 5-flucytosine, no breakpoints are available. NA: Not available.

## DISCUSSION

The emergence of multidrug-resistant *Candida auris* having a high propensity to cause nosocomial transmission render the urgency to develop a novel and effective decolonization strategy. Herein, we demonstrated excellent in vitro fungicidal and/or fungistatic activity of natural antiseptics, including manuka honey, manuka oil and tea tree oil, and their commercial products, against *C. auris*. This suggests the current decolonization strategy can be much improved by combining these well-tolerated, natural antiseptics. A clinical trial on *C. auris* decolonization using manuka body wash, manuka mouthwash, tea tree oil, povidone-iodine, and nystatin oral suspension in 41 patients achieved about 60% success rate and no adverse effects were reported (*Luk*, 2023, unpublished data).

In the present study, the local *C. auris* isolates were highly resistant to both fluconazole (MIC_50_ of 256 µg/mL) and amphotericin B (MIC_50_ of 2 µg/mL). Genomic and proteomic analyses revealed mutation of the fluconazole target enzyme (cytochrome 450 lanosterol 14α-demethylase) [10] and the role of extracellular vesicles in the development of amphotericin B resistance [40].

Our results confirmed the high susceptibility of *C. auris* to 4% chlorhexidine (MFC 0.004%), 10% povidone iodine (MFC 0.312%) and 33.3 μg/mL (100,000 units/mL) nystatin (MFC 9.77 μg/mL) at concentrations used for clinical indications [17, 41]. Nonetheless, the MICs an MFCs of chlorhexidine against *C. auris* strains were higher than those against other *Candida* species. Similar findings were reported by *Abdolrasouli et al* [17], showing that their *C. auris* outbreak strains were less susceptible to chlorhexidine than other *Candida* species though the testing methods were different and the mechanism is currently unknown. Likewise, our clinical *C. auris* strains were highly susceptible to tea tree oil [MFC 1.25% (v/v)] and manuka oil [MFC 0.63% (v/v)]; their fungicidal effects were comparable to synthetic antiseptics, including chlorhexidine, povidone-iodine and nystatin. Our findings are consistent with a previous report of potent antifungal activity of tea tree oil against *C. auris* isolates (MFC 1.56% (w/v)) [20]. Tea tree oil and its main component, terpinene-4-ol, have been shown to cause damage to the fungal cell wall and cytoplasmic membrane thereby causing cellular dysfunctions and death though the exact mechanism is unknown [42–43]. Interestingly, The Body Shop^®^ formulation contains a diluted content of 15% tea tree oil but a lower MIC_90_ of 0.004% (v/v) was observed compared to that of 100% tea tree oil [0.31% (v/v)]. This enhanced inhibition may be attributed to the synergistic or additive interactions of the supplementary ingredients, including denatured alcohol, *Calophyllum inophyllum* seed oil, and *Leptospermum petersonii* oil, which were shown to have antimicrobial properties [44–45]. *Moore et al.* also showed the addition of isopropyl alcohol to disinfectant formulation enhanced the killing of *C. auris* [13]. On the other hand, the antifungal properties of manuka oil were less frequently studied for *C. auris* specifically. *Parker et al.* demonstrated significant inhibitory effects of manuka oil on *C. auris* AR0381 (clade II) and AR0385 (clade IV), with MIC values less than 1.0% [46], of which the findings agree well with ours. More importantly, the low fungicidal concentrations of tea tree oil and manuka oil against *C. auris* are considered safe and tolerable for dermal use. An in vitro skin model proved manuka oil is non-irritative, with cell viability unaffected after treatment with an undiluted sample [47]. Another clinical trial also demonstrated that daily use of 5% tea tree oil shampoo was well tolerated in the treatment for dandruff [48]. The additional property of enhanced skin permeation of essential oils [49] further supports the role of essential oils in eliminating *C. auris* colonization.

Regarding manuka honey, *C. auris* isolates were modestly inhibited (1.5 log reduction in CFU) at a concentration of 25%. The growth inhibition was presented in a dose-dependent manner. This accords with several studies showing fungistatic effect of manuka honey against *Candida* species. [23, 50]. Manuka honey belongs to the “non-peroxide-based” honey group, and its main antimicrobial action is contributed by methylglyoxal (MGO) [51], which is converted non-enzymatically from the dihydroxyacetone present in the nectar of the flowers of *Leptospermum scoparium*. The characteristically high level of MGO in manuka honey makes it a unique marker for authentication. In this study, manuka honey with greater Unique Manuka Factor (UMF), which is correlated with the MGO and total phenol components [52], resulted in greater CFU reduction of *C. auris*. Increasing the concentration of honey from 25% to 50% led to an average 3-log reduction in the CFU of *C. auris* (p < 0.05). Although the inhibitory effect of manuka honey against *C. auris* is modest, honey is seldom applied topically in a clinical setting. Alternatively, honey can act as an important ingredient in the manufacturing of antiseptic products. Of note, the *C. auris* JCM 15448 (clade II) had a much lower MIC (0.2% v/v) towards manuka honey. Similar phenomenon was also observed for manuka oil, tea tree body wash and Medihoney^TM^ wound gel. Comparative genomic analysis revealed that multiple genes involving in cell wall proteins and stress response-related functions were absent in the Japanese *C. auris* strains, including JCM 15448, while a more recent phenotypic analysis also supported clade-specific phenotypic profiles of *C. auris* with clade II having the lowest tolerance to osmotic stress and chemicals [53–54]. These genomic structural variations and metabolic profiles may explain the differences in antifungal resistance, environmental survival, and virulence among different *C. auris* clades. Nevertheless, *de* Groot T *et al.* did not report any difference in antifungal activity of medical grade honey among *C. auris* clades [55]. Studies including larger numbers of *C. auris* isolates of different clades should determine whether susceptibility to natural antiseptics is clade specific.

As expected, commercial products having the active ingredients of natural antiseptics showed promising antifungal effect in vitro against *C. auris*. Manuka body wash and mouthwash were fungicidal against *C. auris* at low concentrations. Product labels of manuka body wash and mouthwash disclose one of the ingredients being 525+ MGO, which is equivalent to UMF 15+ manuka honey. Besides, the aforementioned products contain tea tree and manuka oil with antifungal effects demonstrated in our study. In contrast, Medihoney^TM^ antibacterial wound gel containing 80% medical grade manuka honey and The Body Shop^®^ tea tree body wash were not fungicidal against *C. auris* at concentrations tested though they caused a 6-log reduction in *C. auris* CFU. In addition, the anti-inflammatory and anti-oxidant properties of manuka honey [28] confer the commercial hygiene products as prominent options in the development of an efficacious and tolerable decolonization regimen.

In vitro antifungal susceptibility testing was performed by broth microdilution assay in this study, which is traditionally the gold standard. The MIC and MFC endpoints obtained from broth dilution allows comparison with conventional antifungal agents, for which their clinical correlation has been established; and comparison among different antiseptics or *Candida* species when MIC or MFC data are available. This is particularly useful for evaluating novel antifungal agents for an emerging pathogen. In particular, only limited data of in vitro antifungal efficacy of manuka honey and essential oils are currently available for *Candida* species, including *C. auris*. Our MIC and MFC findings of natural antiseptics were consistent with previous studies using broth dilution assays [20, 23, 46, 50]. Nevertheless, this testing method has some limitations. Contact time exposure is not taken into account during in vitro testing. The exposure time of disinfectants or topical antiseptics in use is often much shorter. Inhibitory and fungicidal endpoints of antiseptics using broth microdilution at 24 hours might therefore overestimate the clinical effectiveness. Alternatively, a time-dependent quantitative assay seems to reflect more about the real-life situation. *Moore et al.* has comprehensively described how to translate the contact time and dilution considerations of antiseptics from clinical use to in vitro testing for *C. auris* [13]. The antimicrobial wash, chlorhexidine 4%, was diluted with water to produce a 2% solution in order to simulate the bath process. When tested with contact time ≤ 2 min, 2% chlorhexidine without isopropyl alcohol failed to eliminate *C. auris*, which was in contrast to the susceptibility demonstrated by broth microdilution after overnight incubation. This highlights the importance of time exposure in the evaluation of antiseptics with regard to the actual use in clinical settings. Moreover, the contact time should be well controlled by neutralizing the antimicrobial activity in vitro after the exposure. *Sutton et al.* has proposed the criteria for identifying the appropriate neutralizing substances, which are specific to the antimicrobial agents and the organisms challenged [56]. Interestingly, a previous study used glyoxalase to neutralize MGO for its main antimicrobial effects in manuka honey. MGO-neutralized manuka honey retained activity against various organisms, indicating the presence of other antimicrobial constituents besides MGO [57]. In addition, most neutralization tests were performed on disinfectants against bacterial organisms and how these results could be applied to fungal organisms was not known. Similarly, there is no universally-agreed neutralizing agent for most natural antiseptics. Alternative approach using membrane filtration or dilution is also unproven on natural antiseptics. Future studies on the active constituents of natural antiseptics will facilitate the validation of appropriate neutralization tests and development of time-dependent assays.

There are some limitations in this study. First, extraction and evaluation of the bioactive chemical markers in the natural antiseptics were not performed. The specific quantities of the active ingredients were often not disclosed in the product labels. The antifungal effects could be contributed by ingredients other than the natural antiseptics we tested individually. We could only tell the overall in vitro antifungal efficacy of the antiseptic products instead of each constituent. Besides, all our *C. auris* isolates belonged to clade I and the results of this study may not be applicable to isolates of other clades. We had included *C. auris* JCM 15448 (clade II/East Asia), ATCC MYA-5002 (clade III/South Africa), and ATCC MYA-5003 (clade IV/South America) in our testing, and found JCM 15448 had significantly lower MIC and/or MFC values towards several natural antiseptics and their commercial products. Moreover, the effect of antifungal drug resistance on the antiseptics could not be studied due to the low variability of the resistance pattern of our *C. auris* isolates. Nonetheless, these antiseptic products had an overall high in vitro antifungal efficacy. Future development of an appropriate time-dependent assay can provide more evidence to support their application as topical or decolonization therapies for *C. auris*.

## CONCLUSION

*C. auris* are highly susceptible to natural antiseptics including tea tree oil, manuka oil and manuka honey, and their commercial hygiene products in the forms of body wash, mouthwash, and wound gel. The promising in vitro results provide insight into the development of novel and effective *C. auris* decolonization regimen. Future studies should demonstrate whether these natural antiseptics are effective in clinical settings.

## ACKNOWLEDGEMENT

The funding of this work was supported by the Clinical Research Centre Mini Research Grant 2023 (2023-CRC-MRG-02), Kowloon West Cluster, Hospital Authority, Hong Kong. We wish to thank the hospital infection control teams in the KW cluster for their efforts in *C. auris* control and the microbiology laboratory staff of Princess Margaret Hospital for the antifungal testing.

## DATA AVAILABILITY

All data are incorporated into the article and its online supplementary material.

