## Supplementary material for "Antifungal efficacy of natural antiseptic products against *Candida auris*"

**S1:** Natural and commercial tested products with ingredients as per label disclosure

| **Natural and commercial tested products** | **Major antiseptic ingredient(s)** | **Other ingredient(s)** |
|---|---|---|
| Manuka oil (Melora®) | Manuka (Leptospermum scoparium) branch/leaf oil | |
| Manuka oil (Manuka Lab®) | Manuka (Leptospermum scoparium) branch/leaf oil | |
| Manuka hydrosol (Bioactive®) | Manuka (Leptpspermum scoparium) branch/flower/leaf water | |
| Tea tree oil (Kiwi Manuka®) | Tea tree (Melaleuca alternifolia) leaf oil | |
| Tea tree oil (The Body Shop®) | Tea tree (Melaleuca alternifolia) leaf oil | Aqua, denatured alcohol, polysorbate 60, limonene, Calophyllum Inophyllum seed oil, Leptospermum Petersonii oil, citral, citronellol, denatonium benzoate and tocopherol |
| Triple action Manuka body wash (Manuka Lab®) | Manuka honey, manuka (Leptospermum scoparium) branch/leaf oil, tea tree (Melaleuca alternifolia) leaf oil | Aqua, ammonium lauryl sulfate, cocamidopropyl betaine, lauryl glucoside, citric acid, capryl glucoside, glycerin, sodium benzoate, potassium sorbate, sodium phylate, Hordeum Vulgare leaf extract, Triticum Aestivurn leaf extract |
| Triple action Manuka mouthwash (Manuka Lab®) | Manuka honey, manuka (Leptospermum scoparium) branch/leaf oil, tea tree (Melaleuca alternifolia) leaf oil | Aqua, sorbitol, polysorbate 20, glycerin, Mentha Piperita oil, Barbadensis Leal juice powder, Triticum Aestivum leaf extract, Hordeum Vulgare leaf extract, menthol, xylitol, caramel, phenoxyethanol, ethylexylglycerin, Capsicum Annuum fruit extract, limonene |
| Tea Tree Skin Clearing body wash (The Body Shop®) | Tea tree (Melaleuca alternifolia) leaf oil | Aqua, sodium laureth sulfate, sodium chloride, peg-40 hydrogenated castor oil, polysorbate 20, cocamidopropyl betaine, peg-120 methyl glucose dioleate, phenoxyethanol, glycerin, sodium benzoate, Calophyllum Inophyllum seed oil, citric acid, butyl methoxydibenzoylmethane, disodium EDTA, limonene, sodium hydroxide, Leptospermum Petersonii oil, tocopherol, caramel |
| Medihoney™ antibacterial wound gel (Comvita®) | Manuka honey | Myristyl myristate, capryl glucoside |

**S2:** Pairwise comparisons of log CFU reduction of *C. auris* isolates by differing concentrations of fungistatic antiseptic products.

| **Reagents** | **p-value** |
|---|---|
| Manuka honey (Comvita®) 15+ | |
| 25% vs. 37.5% | <0.001 |
| 37.5% vs. 50% | <0.001 |
| 25% vs. 50% | <0.001 |
| Manuka honey (Comvita®) 20+ | |
| 25% vs. 37.5% | 0.009 |
| 37.5% vs. 50% | 0.007 |
| 25% vs. 50% | <0.001 |
| Manuka honey (Kiwi Manuka®) 15+ | |
| 25% vs. 37.5% | 0.009 |
| 37.5% vs. 50% | 0.007 |
| 25% vs. 50% | <0.001 |
| Manuka honey (Kiwi Manuka®) 20+ | |
| 25% vs. 37.5% | <0.001 |
| 37.5% vs. 50% | <0.001 |
| 25% vs. 50% | <0.001 |
| Tea tree body wash (The Body Shop®) | |
| 25% vs. 37.5% | 0.003 |
| 37.5% vs. 50% | 0.061 |
| 25% vs. 50% | <0.001 |
